# *Ageratina adenophora* and *Lantana camara* in Kailash Sacred Landscape, India: Current distribution and future climatic scenarios through modeling

**DOI:** 10.1101/2020.09.14.295899

**Authors:** Alka Chaudhary, Mriganka Shekhar Sarkar, Bhupendra Singh Adhikari, Gopal Singh Rawat

## Abstract

The Himalayan region is one of the global biodiversity hotspots. However, its biodiversity and ecosystems are threatened due to abiotic and biotic drivers. One of the major biotic threats to biodiversity in this region is the rapid spread of invasive alien species (IAS). Natural forests and grasslands are increasingly getting infested by IAS affecting regeneration of native species and decline in availability of bio-resources. Assessing the current status of IAS and prediction of their future spread would be vital for evolving specific species management interventions. Keeping this in view, we conducted an in-depth study on two IASs, viz., *Ageratina adenophora* and *Lantana camara* in the Indian part of Kailash Sacred Landscape (KSL), Western Himalaya. Intensive field surveys were conducted to collect the presence of *A. adenophora* (*n* = 567) and *L. camara* (*n* = 120) along an altitudinal gradient between 300 and 3000 m a.s.l. We performed Principle Component Analysis to nullify the multi-colinearity effects of the environmental predictors following *MaxEnt* species distribution model in the current and future climatic scenarios for both the species. All current and future model precision (i.e. Area Under the Curve; AUC) for both species was higher than 0.81. It is predicted that under the current rate of climate change and higher emission (i.e. RCP8.5 pathway), *A. adenophora* will spread 45.3% more than its current distribution and is likely to reach up to 3029 m a.s.l. Whereas, *L. camara* will spread 29.8% more than its current distribution range and likely to reach up to 3018 m a.s.l. Our results will help in future conservation planning and participatory management of forests and grasslands in the KSL– India.

## Introduction

Invasive alien species (IAS) rank among the top three threats to global biodiversity, the other two being unsustainable harvest of various species from the wild and habitat degradation and loss (1). Invasive species, coupled with unsustainable resource use and climate change have seriously affected livelihoods of millions of people in south and south-east Asia (2). Rapid spread of invasive alien species affects natural habitats, regeneration potential of native species and affect productivity of forests and grassland habitats (3). Spread of alien species is particularly challenging in terrestrial ecosystems across the world (4, 5, 6). Changes in climatic conditions and land use practices favour introduction and spread of IAS in most parts of the world (7, 8, 9).Therefore, it is vital to assess the current extent and future scenarios of invasion for various IAS (10,11,12).

The Himalayan region, one of the global biodiversity hotspots, is quite vulnerable due to climate change and other abiotic and biotic drivers of change including rapid spread of IAS. Some of the most obnoxious IAS in this region include *Lantana camara, Mikania micrantha, Chromolaena odorata*, and *Ageratina eupatoria* (13). During the past few decades increasing anthropogenic pressures such as unabated linear infrastructure development, uncontrolled tourism, livestock grazing, agricultural expansion and extraction of non-timber forest products (NTFPs) have led to degradation of wildlife habitats and spread of a large number of invasive species (14). It is estimated that the Indian Himalayan region has over 600 IAS species and nearly 50% of them are said to have escaped from agriculture and accidental introduction (12). One of the major causes of spread of IAS in the Himalayan region is climate change (15). It has been observed that plant species of higher elevation are projected to shift higher, due to which a few IAS previously limited to the lower elevations are now shifting towards the higher altitudes in the Himalaya (15, 16).

As a signatory to Convention on Biological Diversity, India has set its National Target in alignment with Aichi Biodiversity Targets. Accordingly, India’s Biodiversity Strategy and Action Plan aims to identify the priority species of IAS as well as their invasion pathways so that appropriate management strategies could be formulated. It is realized that a very few baseline studies have been conducted on the current extent and spread of even common species in the Himalayan region. Anthropogenic pressures coupled with climate change makes complex the future prediction of species spread. Several authors e.g., (15, 17-21); have opined that the IAS which were earlier limited to the lower areas are likely to shift towards the higher elevations in the Himalaya. This calls for validation of earlier models as well as multi-locational studies. In this paper, we present the findings of a case study on two IAS, viz., *A. adenophora* and *L. camara* at a landscape level within Indian part of Kailash Sacred Landscape, hereafter referred as Kailash Sacred Landscape (KSL) –India. Major objectives of the study were to predict potential current spread of these species in KSL–India and predict their invasion in future climate change scenarios.

### Study Area

The study was carried out in KSL – India that is located between 29°26’35” to 30°35’13” N and 80°01’24” to 81°02’44” E. It is characterized by heterogeneous landscape, wide altitudinal range (ca. 800 to over 7000 m a.s.l.), diverse topography and rich biodiversity (17). This area is contiguous with far-western Nepal and Tibet Autonomous Region (TAR) of China. The study area is spread over 7,120 km2 area (Figure 1).

**Figure 1:**
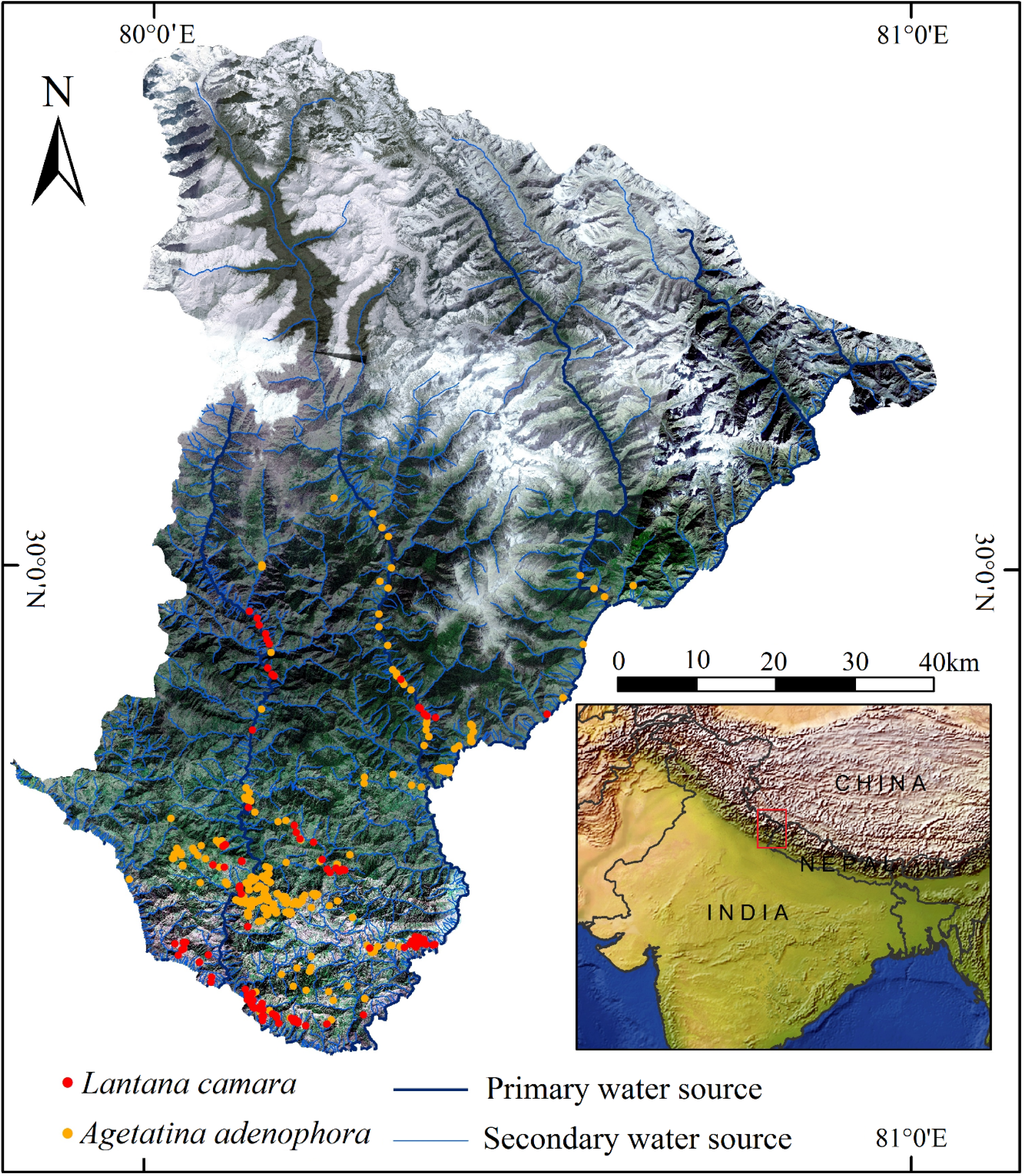
Geographical location of the study area with sampling locations of *Lantana camara* and *Ageratina adenophora*.

KSL – India has the predominance of diverse natural ecosystems from grasslands to moist sub-tropical broadleaved to temperate oak forests, sub-alpine conifers, high altitude birch forests, while extensive alpine pastures occur in areas >3000 to 4000 m a.s.l beside agro-ecosystems in lower reaches (22-23). The diverse habitats cater to numerous indigenous flora and fauna. Forests in KSL – India fall into two categories - Protected Areas (PAs) and non-protected areas. KSL – India includes only one legally designated PA (*i*.*e*. Askot Wildlife Sanctuary) which provides important ecosystem services for the region. However, the non-PA category of forests includes reserve forest and protected forests under the control of the State Forest Department of Uttarakhand managed for an extended period for production of timber and non-timber forest products (NTFPs). KSL – India also includes highly interspersed small forest fragments falling under two broad categories, *i*.*e*. civil/soyam forests and community forests are under the control of Revenue Department of the State Government. The latter category of community forests, for nearly nine decades or so, has been managed by the local communities and is referred to as ‘*Community Forests*’ or ‘*Van Panchayat Forests*’. This community forest serves as the primary source of livelihood to locals. The KSL – India has experienced a considerable change during recent decades in terms of land use and land cover, along with the development of infrastructure and patterns of natural resource used by the local communities (23-24).

## Methods

### Species presence

After the extensive vegetation survey and consultation with experts, two most widely spread IAS, i.e. *Ageratina adenophora* (Asteraceae) and *Lantana camara* (Verbenaceae) were selected for the present study. *A. adenophora* is native of Central America (25) and distributed worldwide (26). *L. camara*, native to Tropical America (27) is considered to be one of the most troublesome invasive species worldwide (28). *A. Adenophora*, a perennial, herb produces a large number of tiny seeds which are dispersed by wind and human activities (29). Its ubiquitous property to invade diverse habitats, such as roadsides, riverbanks, forest edges, crop fields, wastelands, and rubbish dump edges makes it fit to invade various ecosystems. *L. camara* is one of the most widely distributed IAS in India (30), resistant to fire and can regrow if burnt (31). It reproduces primarily by seeds as well as through coppice and prefers to grow in degraded habitats.

Though, the species selected for the study are known to negatively affect the wildlife habitats and native vegetation, they are also known to a have a few minor use to local communities. Leaves of *A. adenophora* are used for cattle bedding (32), or its leaf paste is applied to cuts and wounds (33). The wood of *L. camara* is used as fuel (34), source of food for birds (35-37) and nectar for butterflies and moths (38) as it flowers and fruits throughout the year.

### Presence record

We recorded *A. adenophora* and *L. camara* presence between 300 and 2500 m a.s.l covering a wide range of habitats including agricultural lands, grasslands, fallow lands and forests, proximity to major road networks and naturally occurring drainages in different parts of the landscape during 2014 to 2017. A total of 567 and 120 presence locations for *A. adenophora* and *L. camara* respectively, were recorded using handheld Garmin N72 GPS. Of these, 80% locations were used for probability distribution modeling and the rest 20% were used to evaluate the model performance.

### Selection of environmental and bioclimatic variables

We used two-time periods (“current” and “2050”) environmental data to model the *A. adenophora* and *L. camara* current and future distribution in KSL – India. A total of 22 environmental variables were used for probability distribution modeling in each time periods. We retrieved standard 19 bioclimatic variables for WorldClim version 2 (39). These are the average for the years 1970-2000 with a spatial resolution of 30 seconds (∼1 km2). These variables are considered as current bioclimatic variables. We also retrieved 19 bioclimatic variables with a spatial resolution of 30 seconds (∼1 km2) for the year 2050 (average for 2041-2060). These are the climate projections as per the Intergovernmental Panel on Climate Change (IPCC) 5th assessment report from Global Climate Models (GCMs) for one Representative Concentration Pathways (RCPs). The data were downscaled and calibrated using WorldClim 1.4 as baseline ‘current’ climate (39, 40). These bioclimatic data are also the most recent GCM climate projections that have already been considered in the IPCC fifth assessment report.

We also considered four major static topographic factors for both the “current” and “2050” time frame. These are elevation (ASTER, Digital elevation data from Advanced Space-borne Thermal Emission and Reflection Radiometer), slope, aspect, Euclidian distance from water channels and Euclidian distance from major roads. These variables were assumed to be constant across the entire study period. The details of all the variables used in this study are provided in Appendix 1.

### Pair-wise Pearson correlations and Principal Component Analysis

We checked for multi co-linearity with pair-wise correlations for both “current” and “2050” environmental data sets in order to avoid spurious model calibrations (41). It is common a practice to retain only predictors with pair-wise correlation < |0.7| (42). To nullify the correlation structure of the variables and inter-relationship among them, we implemented the Principal Component Analysis (43-44). The PCA reduces the number of orthogonal predictors each of which is the linear combination of 24 original environmental variables. All the exploratory variables were subjected to PCA to extract the factors of significant contribution using *prcomp* function and predicting function of R version 3.3.3 (45). Furthermore, we obtained 24 Principal Components (PCs) in raster format. For both “current” and “2050” data sets, the first six PCs accounted for 99% of the variability and were chosen for further multivariate species distribution modeling (Table 1).

**Table 1.** Representation of the first seven principal components derived from all 24 original environmental variables along with the proportion of variance in environmental variables explained by each principal component and their cumulative proportion of variance.

| Principal components | The “current” year |  | Year 2050 |  |
| --- | --- | --- | --- | --- |
|  | Proportion of Variance | Cumulative Proportion | Proportion of Variance | Cumulative Proportion |
| PC1 | 0.98890 | 0.98890 | 0.98880 | 0.98880 |
| PC2 | 0.00810 | 0.99700 | 0.00810 | 0.99700 |
| PC3 | 0.00276 | 0.99979 | 0.00277 | 0.99973 |
| PC4 | 0.00013 | 0.99992 | 0.00015 | 0.99988 |
| PC5 | 0.00007 | 0.99998 | 0.00008 | 0.99996 |
| PC6 | 0.00001 | 0.99999 | 0.00003 | 0.99999 |
| PC7 | 0.00000 | 1.00000 | 0.00000 | 1.00000 |

### MaxEnt species distribution modeling

Maximum Entropy (*MaxEnt*) distribution modeling was implied to identify the current and future invasion for both, *A. adenophora* and *L. camara* in KSL – India. *MaxEnt* is a machine learning algorithm which is widely used for species distribution modeling that uses presence-only data (46-47) to predict potential distribution of a particular species in the current as well as future climatic scenarios (48-49).

The occurrence data from the field was collected for both *A. adenophora* (*n* = 567) and *L. camara* (*n* = 120). One thousand background points were drawn randomly across the KSL – India by considering pseudo-absence of both species. We generated four main *MaxEnt* models (two for each species) using the six PCA derived explanatory variables for both ‘current’ and ‘2050’ year. We implemented k-fold cross-validation (50) to build each main *MaxEnt* model. Firstly, we used standard k-fold cross-validation in our randomly partitioned approach. In our k-fold cross-validation approach, the occurrence localities were divided randomly into 10 bins, where each bin constitutes an equal proportion of sample size (51-52). Then, each model was developed iteratively, using (k − 1) bins for model calibration in each iteration, with the remaining fold retained for evaluation. This is repeated until all the folds were utilized at-least once for model evaluation. *MaxEnt* produces a model based on a series of ‘features’, such as a linear, hinge, quadratic, product, threshold and discrete; we used all except for discrete, as all of our explanatory variables were continuous (53). In the ‘feature’ functions, we set beta multiplier as 0.5 (medium) that affect the smoothness to the model output. We have opted for clamping function in each model as the use of the function is strongly suggested for projecting future species distributions by both Elith et al. (54) and Webber et al. (55).

As we set the models *via* k-fold cross validation (*k* = 10), for each dataset, we used the respective evaluation localities to calculate Area Under the Curve (AUC) of the Receiver Operation Characteristic (ROC), which is a measure of the overall discriminatory ability of the model or to precisely evaluate model performance by plotting its model sensitivity graph (56). We averaged those values from each sub-models of each main-model and represented the averages for each of the four main *MaxEnt* models. The AUC score closer to 1 indicates that the model had accurately predicted the habitat, while a value ≤0.5 indicates low accuracy of the model (57).

Binary habitat invasion maps for the ‘current’ and future ‘2050’ were prepared by using the threshold of maximum training sensitivity plus the specificity logistic threshold. The threshold approach of ‘*Maximum training sensitivity + specificity*’ is a more reliable, restrictive and conservative approach to understand the potential distribution of a species. This particular threshold approach is one of the popular methods either for presence/absence or presence-only data (58). We used ‘*dismo*’ package in R (59) along with a source file developed by Feng et al. (60) to build all *MaxEnt* models as well as to evaluate their performance.

### Invasion across the elevation gradient

To check the future spread of *A. adenophora* and *L. camara*, we extracted all elevational range values from a 30 m resolution digital elevation model (ASTER) using the threshold polygons generated from *MaxEnt* main models. Elevational gradients fall within the distribution ranges were re-classified into multiple classes considering 100 m a.s.l intervals. Area invaded by both species in each elevational class were compared between current and 2050 year using paired sample t - test. The distributions of elevational class vs. invaded areas of both *A. adenophora* and *L. camara* were plotted separately for the current year and 2050 to visually asses their spread pattern across the elevational classes.

## Results

The pair-wise Pearson correlations were generally higher (r >|0.70|) for most of the variable (Figure 2). We were not be able to use these variables directly in *MaxEnt* as they showed high correlation, and this avoids the overprediction of suitability in *MaxEnt* models. However, PCA transformed the 24 environmental variables (each for ‘current’ and ‘2050’) into a composite set of 24 components that are orthogonal and un-correlated to each other. Six out of 24 components demonstrated the cumulative proportion of variance > 0.99 for both the ‘current’ and the year ‘2050’ (Table 1). The first six PCs each from the ‘current’ and ‘2050’ were represented in Appendices 2-3.

**Figure 2:**
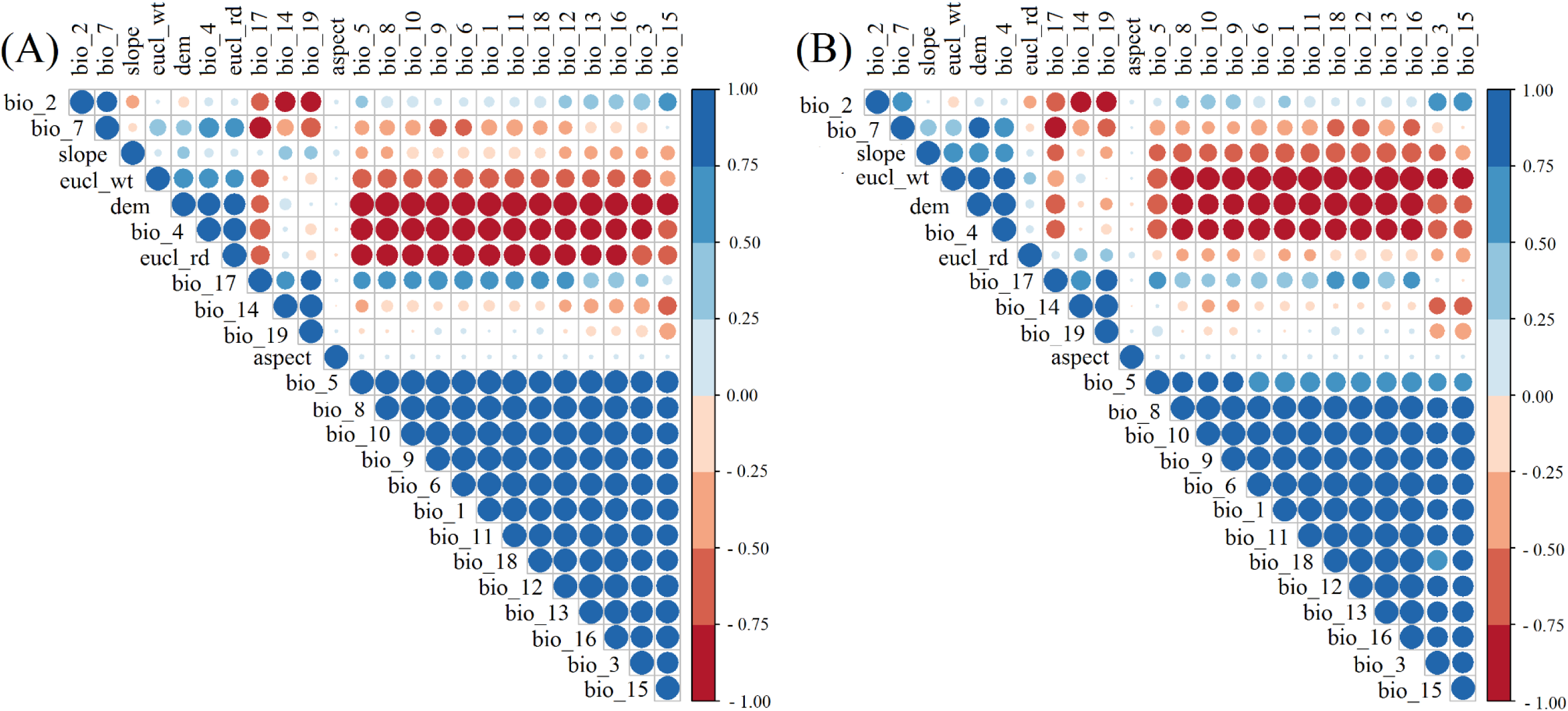
Representation of Pearson correlation matrix for environmental data of the “current” year (A) and year 2050 (B).

Both training and test area under ROC curve (AUC) values were > 0.81 for both species under the ‘current’ and future ‘2050’ climatic conditions (Figure 3, Table 2). The current occurrence locations of *A. adenophora* and *L. camara* where 95.06% and 92.5% respectively falls within their most climatically suitable current distribution ranges. The relative percent contribution of all PCs from both the ‘current’ and the year ‘2050’ are presented in Table 3, and all response curves associated with the above MaxEnt model predictions were given in Appendix 4.

**Table 2.** The area under the receiver operating curve (AUC) score of MaxEnt models for both *Ageratina adenophora* and *Lantana Camara*.

| Species | The “current” year |  | Year 2050 |  |
| --- | --- | --- | --- | --- |
|  | Test AUC | Training AUC | Test AUC | Training AUC |
| *Ageratina Adenophora* | 0.81 | 0.82 | 0.81 | 0.82 |
| *Lantana Camara* | 0.93 | 0.93 | 0.92 | 0.93 |

**Table 3.** Relative contributions of each principal component are used in all the *MaxEnt* models.

|  | Contribution of *Ageratina adenophora* (%) |  | Contribution of *Lantana camara* (%) |  |
| --- | --- | --- | --- | --- |
|  | The “current” year | Year 2050 | The “current” year | Year 2050 |
| PC1 | 83.00 | 86.10 | 62.8 | 65.2 |
| PC2 | 00.00 | 00.10 | 0.1 | 0.2 |
| PC3 | 01.10 | 02.40 | 9.1 | 15.6 |
| PC4 | 04.60 | 04.60 | 2.1 | 1.1 |
| PC5 | 07.30 | 05.40 | 16.7 | 15.8 |
| PC6 | 04.00 | 01.50 | 9.1 | 2.2 |

**Figure 3:**
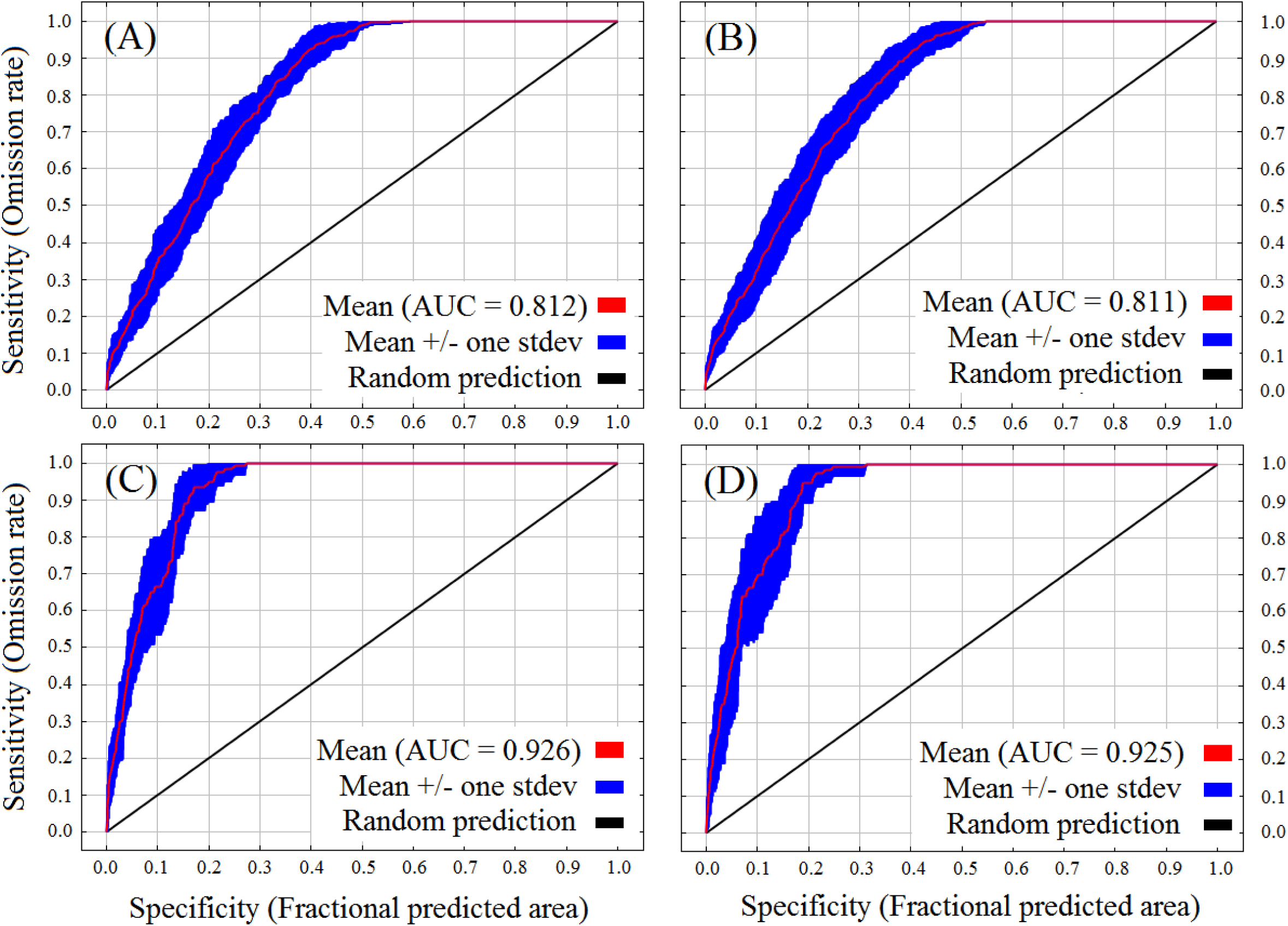
Receiver operating characteristic (ROC) curve for the predicted distribution *MaxEnt* models averaged over the ten replicate runs: *Ageratina adenophora* (the “current” year – A; year 2050 - B) and *Lantana camara* (the “current” year – C; year 2050 - D).

The current and future distributions of the *A. adenophora* and *L. camara* along the elevational gradient of KSL – India are determined by the RCP 8.5 pathway projections for 2050 (Figure 4A-B).

**Figure 4:**
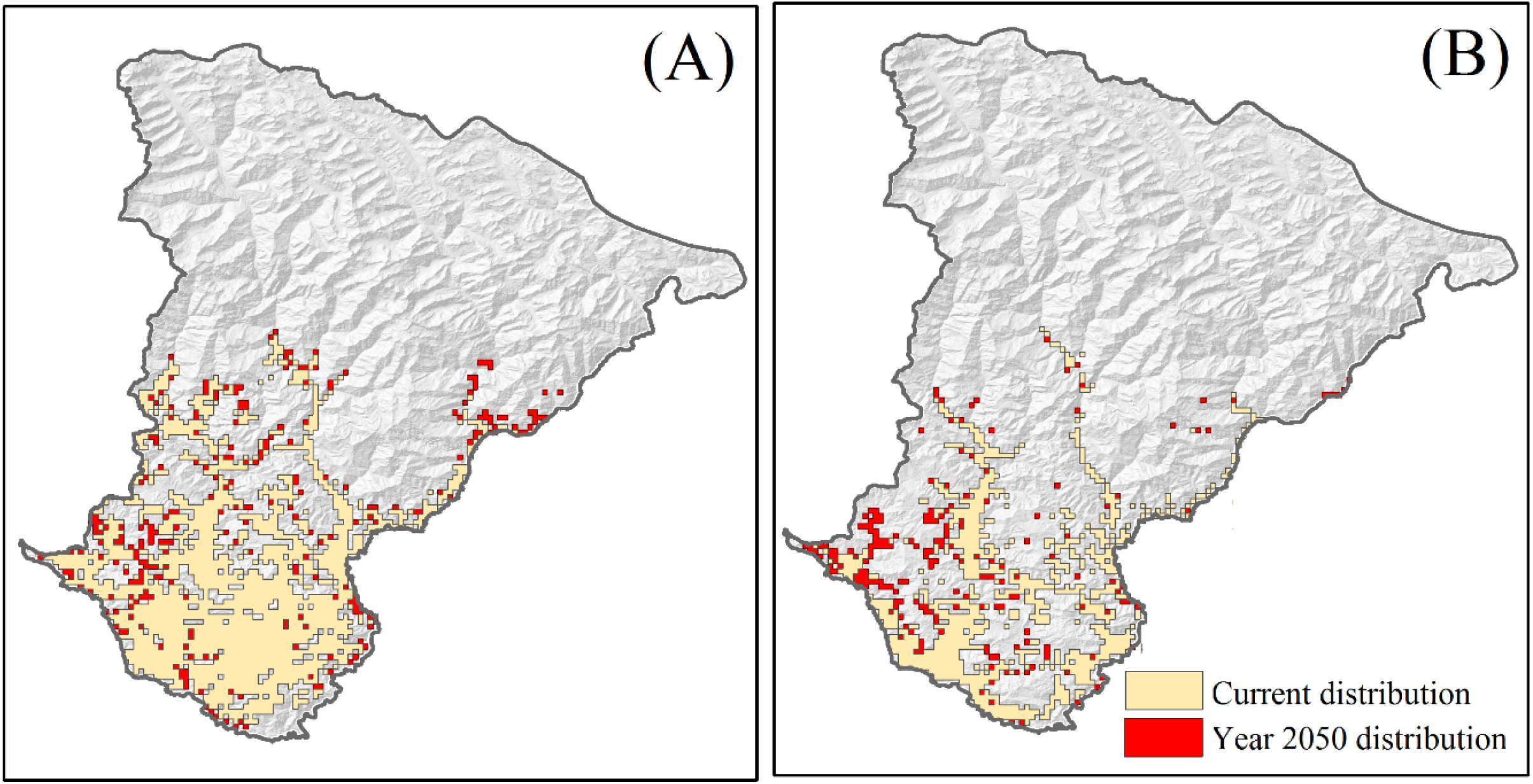
Predicted distribution *MaxEnt* models averaged over the ten replicate runs. Orange and Red polygons represent maximum training sensitivity + specificity logistic threshold for the distribution of (A) *Ageratina adenophora* and (B) *Lantana camara* in the “current” and year 2050.

The current distribution of *A. adenophora* ranges between 212 - 2616 m a.s.l, while *L. camara* distributes mostly below 2045 m a.s.l. The potential area that is currently vulnerable to invasion by *A. adenophora* and *L. camara* extends to 1403.7 km2 and 1053.1 km2, respectively. If the climate change continues to follow the higher emission RCP 8.5 pathway, by 2050, invasion of *A. adenophora* would reach up to 3029 m a.s.l, whereas, *L. camara* will invade up to 3018 m a.s.l. Both *A. adenophora* and *L. camara* will spread across the landscape occupying an area of 2039.7 km2 and 1366.5 km2, respectively. The area invasion by *A. adenophora* (*t* = -7.61, *df* = 28, *p* ≤ 0.05) and *L. camara* (*t* = -2.70, *df* = 28, *p* ≤ 0.05) will significantly increase by year 2050 across all designated elevational class (Figure 5).

**Figure 5:**
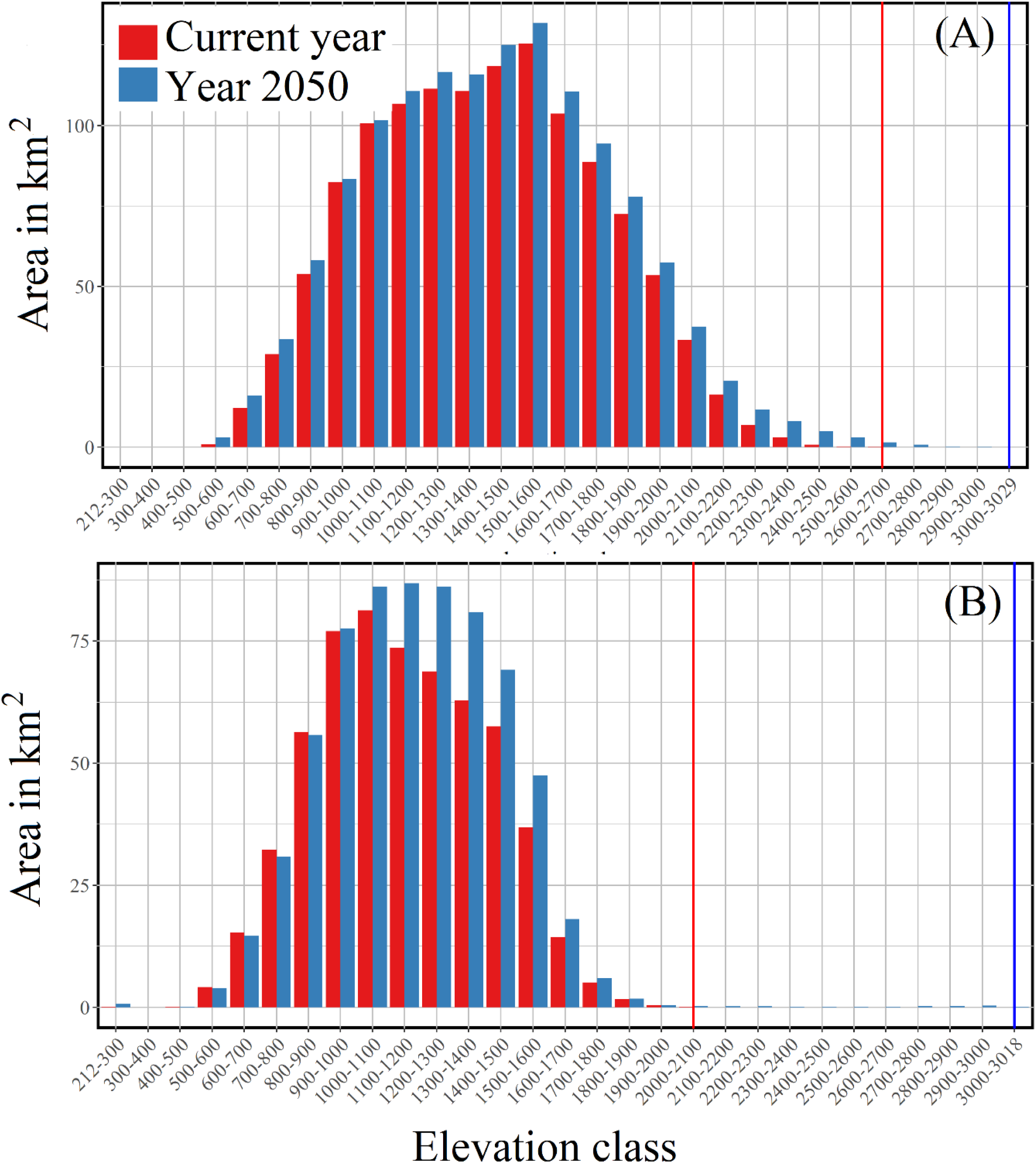
The predicted spread (in Km^2^) of *Ageratina adenophora* (A) and *Lantana camara* (B) in different elevational classes of KSL – India during the current year and the year 2050.

## Discussion

Plant invasion is the emerging area of science that impacts substantial economic and ecological imbalance by altering native plant community composition (61) and depleting native plant species diversity (62) thus affecting ecosystem processes (63-65). The lower and mid ecosystems in the mountains are more vulnerable to invasion by exotic plant species due to the altering climate change and increasing anthropogenic pressure (66). These plant species also exceeded their ranges and many of them have become abundant at higher elevations (67-68). The increasing incidence of invasion in the high-altitude ecosystems (9) poses a significant threat to the native biological diversity of the region and it is expected to expand in the future (21). The use of new and robust methods, especially combining PCA and *MaxEnt* species distribution modeling will reduce the chance of overpredicting species distribution map.

This demonstrated an increasing (both vertical and horizontal) trend of plant invasion in KSL – India. Altering climatic condition and mostly increased temperature have been providing favourable conditions for the invasion and spread of these species in the KSL-India. Joshi et al. (69) already demonstrated that *L. camara* occurrence was recorded widely in dry and exposed slope, forest fringes and deep forests throughout their study sites in the Indian butter tree (*Diploknema butyracea*) forest which is one of the economically, spiritually and culturally significant trees in the region with its great importance for local livelihood. Looking forward, *L. camara* might also affect the regeneration of such economically important trees in the area. Elevation was the most crucial factor in the current and future distribution. It appeared that in future both of these IAS would colonize more vigorously in the landscape where currently they are not present. This invasion of both species is likely to happen if climate change continues to follow the higher emission RCP 8.5 pathway. Furthermore, *L. camara* will spread within a similar altitudinal range where it is currently absent. According to Singh et al. (70), *A. adenophora* prefers north-facing moist slopes along roadsides, forest fringe, cliffs, near water bodies, including agricultural and degraded lands and sub-tropical and warmer temperate landscapes. It also invades natural habitats, Deodar, Banj oak-Chir pine and mixed forests, degraded grassland areas and sometimes invading pure stands of pristine temperate broadleaved oak (*Quercus leucotrichophora*) forest representing climax vegetation between 1000 – 3500 m a.s.l. in KSL – India. Our study also showed a similar trend, as this species will reach 968 m a.s.l higher than its current distribution range.

Our predictive models also depicted high infestation and expansion in the present climatic scenario by *A. adenophora* and *L. camara* in the lower parts of KSL – India in western Himalaya. The MaxEnt model also predicted that the landscape provides more suitable conditions for the infestation of *A. adenophora* as compared to *L. camara* even at its current climatic condition. Both the species will expand their habitats with the increase in temperature and human activity in the landscape. Our study demonstrated similar results as compared to previous studies (71-73), but in a more robust and spatially explicit manner.

Our study has come up with current pattern of distribution and established future scenarios of invasion by two highly obnoxious IAS within KSL – India. The distribution maps generated in this exercise would be of much help in further conservation planning and participatory management of forest / grassland management. The findings of our study can be used in identification of vulnerable areas and important localities which are likely to be infested by these species. Habitat restoration and community-based monitoring at local scale involving Biodiversity Management Committee (BMC) would be one of the logical steps forward.

## Acknowledgments

The authors are thankful to the Director of the Wildlife Institute of India for his support. The study was carried out under Kailash Sacred Landscape Conservation and Development Initiative (KSCDI) funded by ICIMOD, Nepal and National Mission on Himalayan Studies (NMHS) funded by Ministry of Environment, Forests and Climate Change, New Delhi. We would also like to acknowledge our gratitude to the local communities for their help, support, and cooperation during this study.

**Appendix 1.**
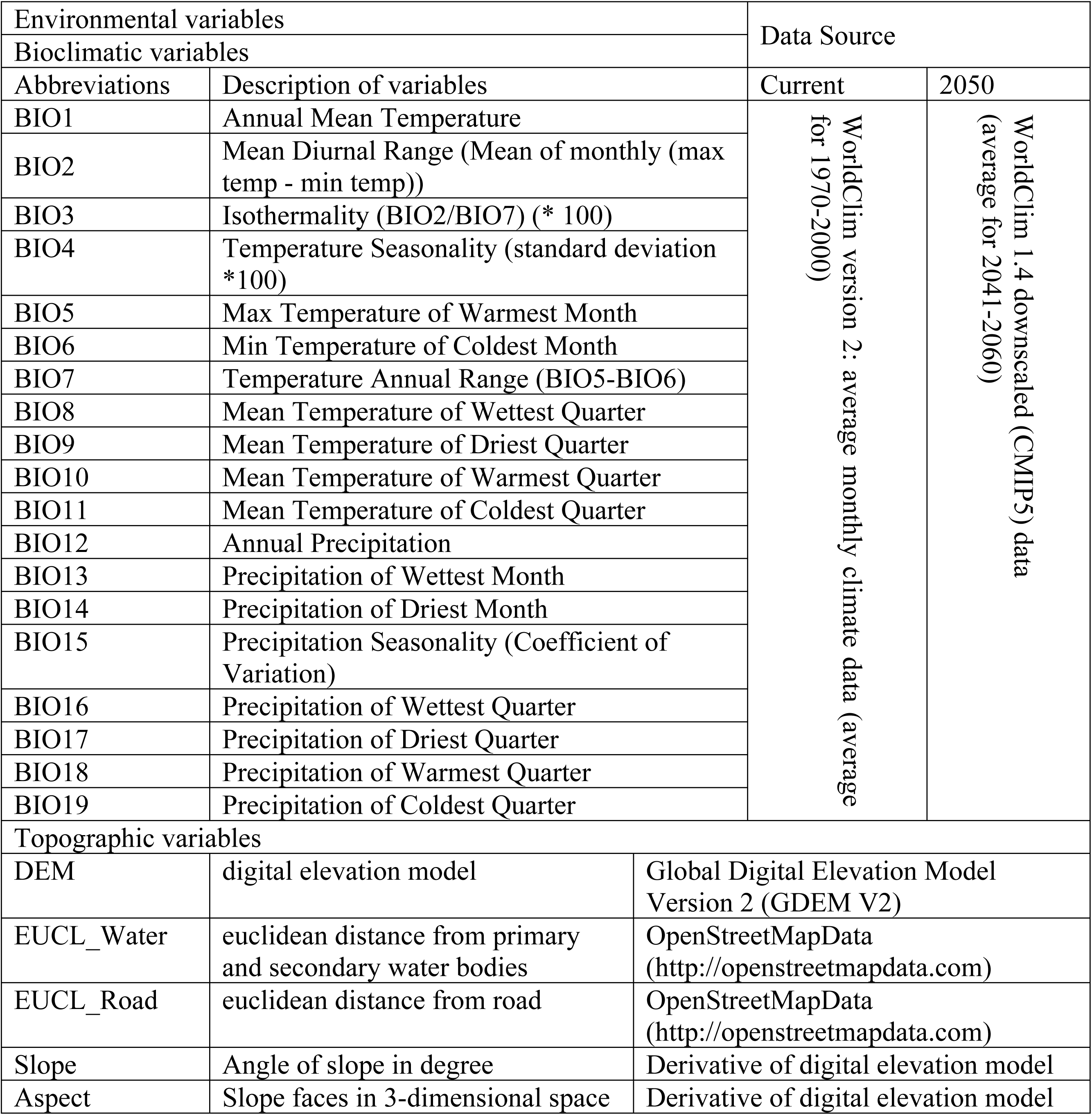
Description of variables of three time periods used to model *Ageratina adenophora* and *Lantana camara* distribution across the KSL-India.

**Appendix 2.**
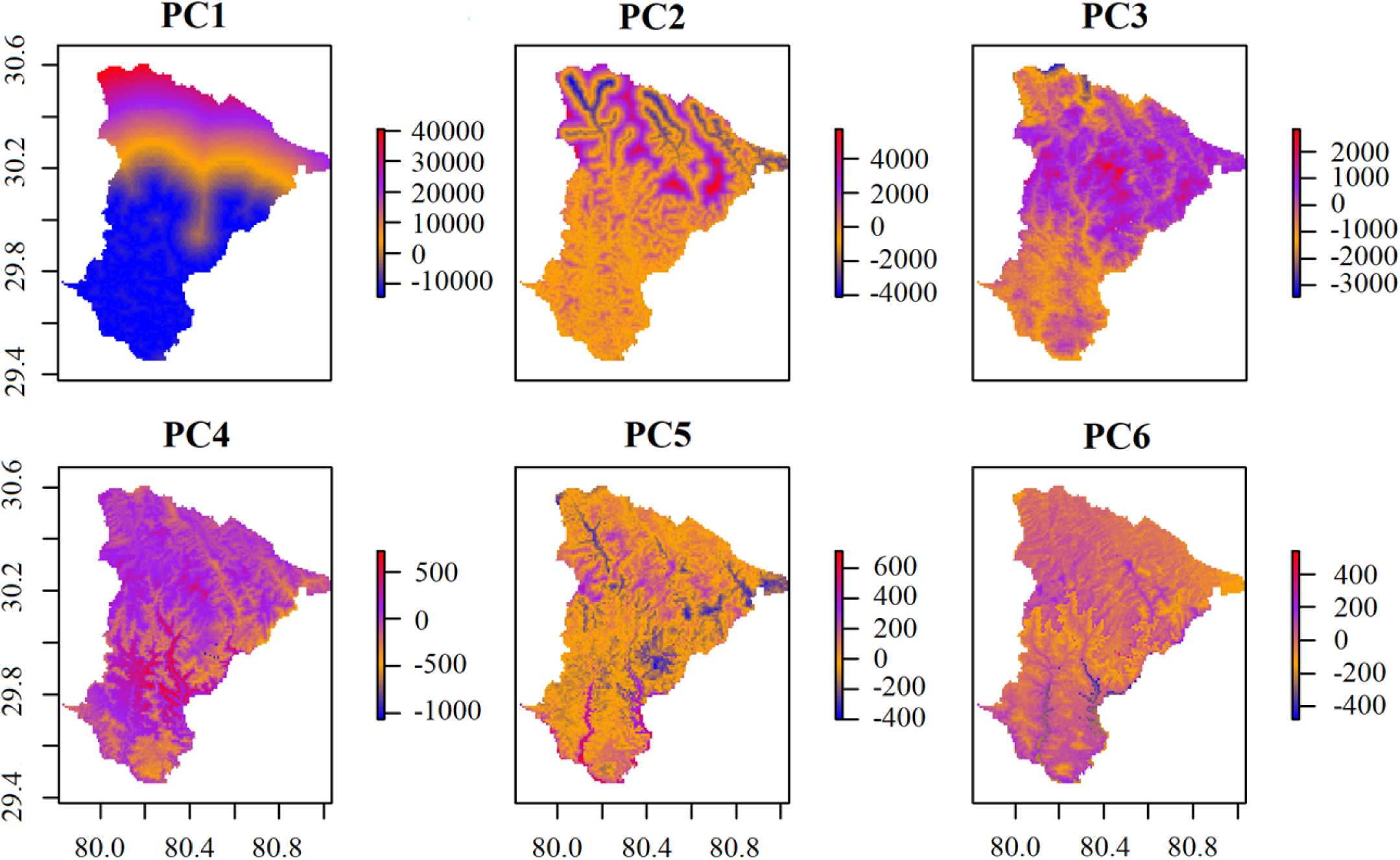
First six principal components as raster layers generated from the 24 environmental layers of current years.

**Appendix 3.**
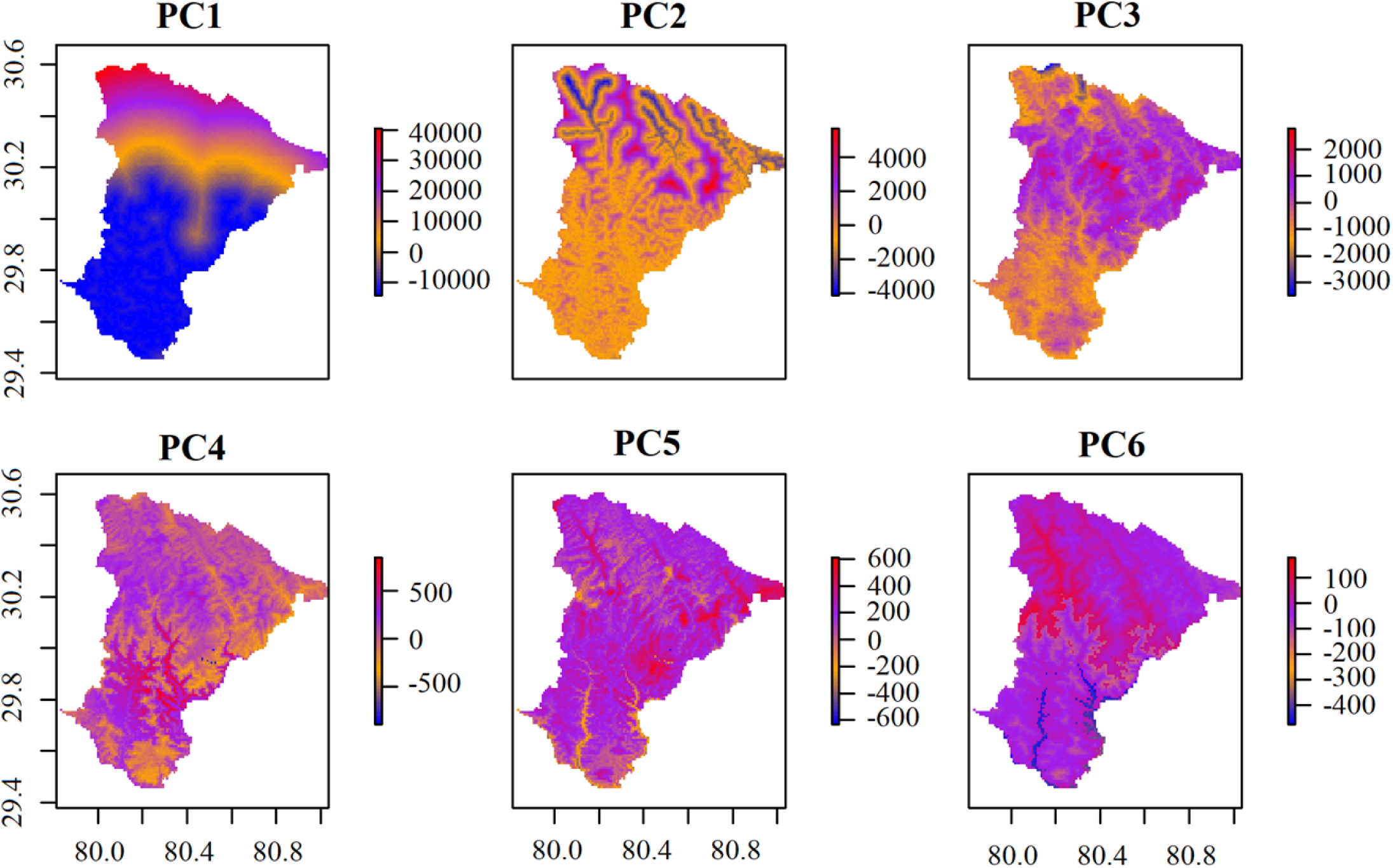
First six principal components as raster layers generated from the 24 environmental layers of 2050 years.

**Appendix 4.**
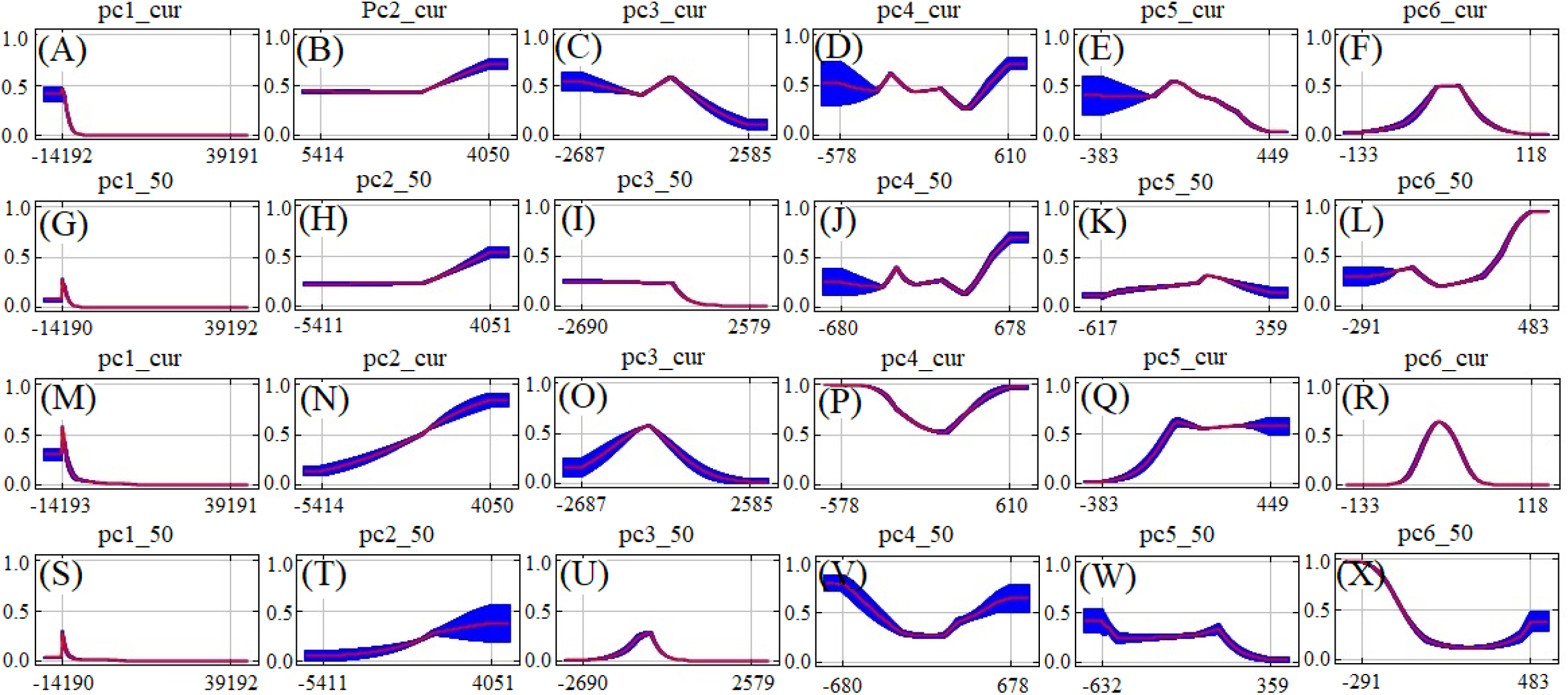
These curves show how each PCs affects the MaxEnt prediction. The curves show how the predicted probability of presence changes as each PC is varied, keeping all other environmental variables at their average sample value. The curves show the mean response of the 10 replicates of each *MaxEnt* runs. (A-F): Response curves of *Ageratina adenophora* for current year, (G-L): Response curves of *Ageratina adenophora* for the year 2050, (M-R): Response curves of *Lantana camara* for current year, (S-X) Response curves of *Lantana camara* for the year 2050.

